# Contextual influences of visual perceptual inferences

**DOI:** 10.1101/2022.09.15.507892

**Authors:** Emily J A-Izzeddin, Jason B Mattingley, William J Harrison

**Affiliations:** Queensland Brain Institute, University of Queensland, Australia; School of Psychology, University of Queensland, Australia

## Abstract

Humans have well-documented priors for many features present in nature that guide visual perception. Despite being putatively grounded in the statistical regularities of the environment, scene priors are frequently violated due to the inherent variability of visual features from one scene to the next. However, these repeated violations do not appreciably challenge visuo-cognitive function, necessitating the broad use of priors in conjunction with context-specific information. We investigated the trade-off between participants’ internal expectations formed from both longer-term priors and those formed from immediate contextual information using a perceptual inference task and naturalistic stimuli. Notably, our task required participants to make perceptual inferences about naturalistic images using their own internal criteria, rather than making comparative judgements. Nonetheless, we show that observers’ performance is well approximated by a model that makes inferences using a prior for low-level image statistics, aggregated over many images. We further show that the dependence on this prior is rapidly re-weighted against contextual information, whether relevant or irrelevant. Our results therefore provide insight into how apparent high-level interpretations of scene appearances follow from the most basic of perceptual processes, which are grounded in the statistics of natural images.

## Introduction

We move through different environments many times each day. Each environment exposes our visual system to a unique combination of features that guide cognition and behaviour (Frazor & Geisler, 2006; Torralba & Oliva, 2003). When entering a new environment, we can readily categorise and identify how its features relate to our current goals (Greene & Oliva, 2009; Walther et al., 2009). Our ability to make such flexible assessments suggests that visual cognition leverages internally stored sets of rules that help to make sense of new sensory information.

The sets of rules that guide visual cognition can be considered as expectations, or priors, for visual features that are common across environments. Priors putatively represent the average of features on any number of dimensions (Series & Seitz, 2013; Summerfield & Egner, 2009). When incoming sensory information deviates from a set of priors, our expectations are violated. However, the inherent variability in image features across environments (Hansen et al., 2003, 2008) does not appreciably challenge our visual or cognitive capacities. While priors may be a useful guiding tool when interpretating our environment, therefore, they must be flexible, allowing functional interpretation of a vast array of possible visual scenes. Indeed, expectations shift depending on context-specific information (Brockmole & Le-Hoa Vo, 2010; Wolfe et al., 2011), suggesting a trade-off between priors that generally apply in the longer term versus those that depend on a specific context.

Visual information can be quantified at any arbitrary level, but in the present study we broadly categorise features as being either low- or high-level. By low-level information, we refer to basic visual features, such as orientation, contrast, or hue. Under the same framework as Neri (2014), we consider high-level information to include features which convey meaningful information (e.g., the arrangements of chairs and a table that imply a dining room setting as opposed to an office). Importantly, two images can be matched on a subset of low-level features, but have these features arranged such that only one conveys meaning (Neri, 2014). Low-level features, such as contrast energy across spatial frequency bands, are relatively stable across contexts, whereas many attributes of high-level features are scene-specific (Harrison, 2022; Torralba & Oliva, 2003), and their respective statistical regularities across scenes influence decision making. For example, cardinally oriented features are overrepresented in natural images (Coppola et al., 1998; Essock et al., 2003; Girshick et al., 2011; Hansen et al., 2003; Hansen & Essock, 2004; Keil & Cristóbal, 2000), leading to well-known biases in perceptual judgments of orientation (Appelle, 1972; Berkley et al., 1975; Campbell et al., 1966; Dakin, 2001; Dakin et al., 2009; Dakin & Watt, 1997; de Gardelle et al., 2010; Emsley, 1925; Girshick et al., 2011; Pratte et al., 2016; Westheimer & Beard, 1998). Similarly, relationships between high-level visual objects can be predicted from scene context, leading to errors in object detection and recognition when objects are positioned at uncommon locations (Bar, 2004; Bar & Ullman, 1993; Biederman et al., 1982). Despite their differences in content and contextual stability, we combine low- and high-level features with great ease to allow a seamless percept of our environment. However, the relative contribution of low- and high-level information to our interpretations of a scene remain largely unknown.

In the present study, we investigated the interplay between relatively long-term priors and context-dependent information in forming visual expectations. To force observers to use and combine previous experience with changing high-level contextual information, we took advantage of the statistical distributions of features present in naturalistic stimuli (David et al., 2004; Harrison, 2022; Olshausen & Field, 1996; Simoncelli & Olshausen, 2001). Targets were randomly oriented natural image patches that participants rotated to appear subjectively “upright” based on their own internal criteria (see Methods for specific task design information). By windowing the targets within a small aperture, we removed large-scale contextual information. In the absence of such high-level information, we anticipated that observers would base their responses on how closely target features match their internal priors for low-level image features alone. In some conditions, the surrounding region of the target was presented briefly prior to target presentation, allowing observers to incorporate contextual information to inform their judgments. Our task, therefore, required participants to align targets with their internal expectations formed from both longer-term priors as well as immediate contextual information, thereby enabling us to disentangle the relative contribution of each.

## Results

### Natural image statistics predict perceptual inferences in the absence of context

Participants rotated natural image target patches to make the targets appear upright (see Fig. 1A-B). Targets contained very little high-level structure that could unambiguously inform responses, as we confirmed in control experiments (described below and in Supplemental Materials). By having participants rotate the targets, they were able to control various low-level image features, such as the relative frequency of orientations present. In Experiment 1, targets were presented without context. If participants responded randomly, their responses would be uniformly distributed. If instead participants used any sort of strategy that depended on the target image features, their responses would be systematically biased by their strategy. For example, if participants were biased by orientation information present in the targets, then we might observe response orientations biased towards particular orientations, such as cardinals.

**Figure 1.**
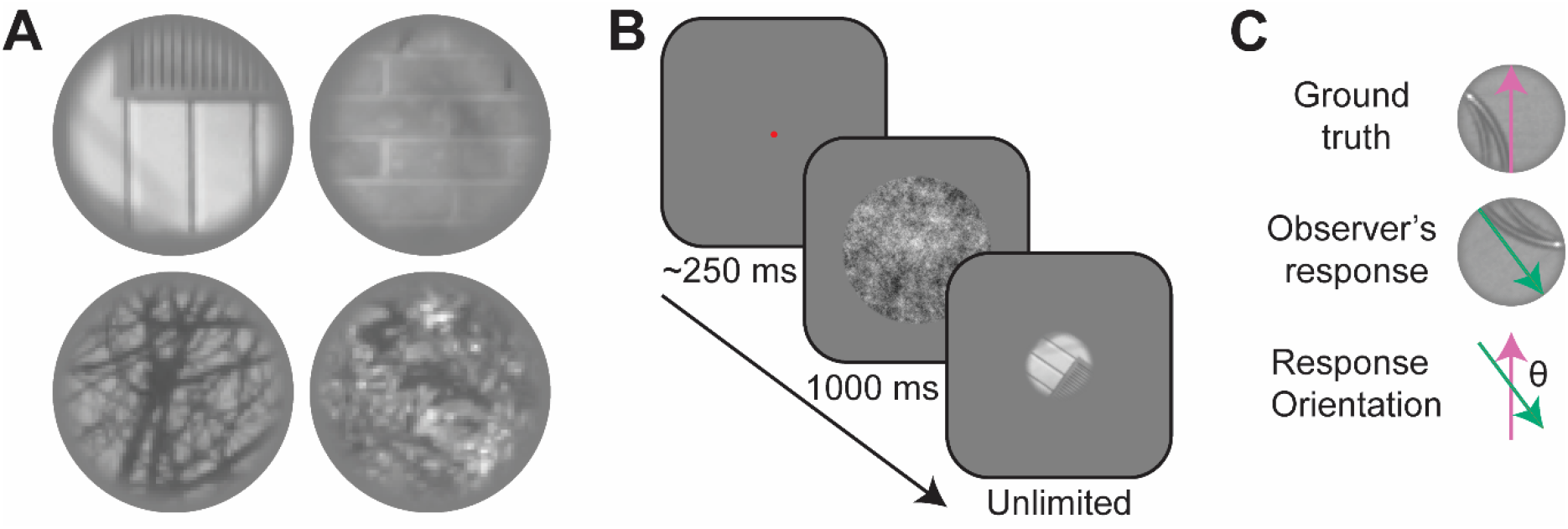
Overview of paradigm. **A)** Example windowed image patches used as targets, shown to participants at random orientations from trial to trial. **B)** Example schematic for a trial in Study 1 (stimuli are not to scale). Participants saw a fixation point, followed by a brief pink noise patch. Subsequently, the target was presented at a random orientation. Participants used the mouse to rotate the target to appear upright and clicked to input their response. **C)** Observers’ judgments were circularly distributed deviations from the known objective vertical axis of the target. Note that the top left target patch in (A) is rotated 90° from upright, demonstrating the inherent ambiguity in the stimuli.

The blue curve in Figure 2A shows the frequency distribution of participants’ responses relative to the objective upright orientation of each target. Participants’ responses have a clear cardinal bias: the most frequent response orientation is centred on 0°, demonstrating participants’ modal response was highly accurate, with smaller peaks at the other cardinal orientations (±90° and ±180°). The multimodal distribution of responses reveals that participants were not simply guessing. Instead, observers must have combined target information with some internal representation of the most appropriate orientation given the features in the image.

**Figure 2.**
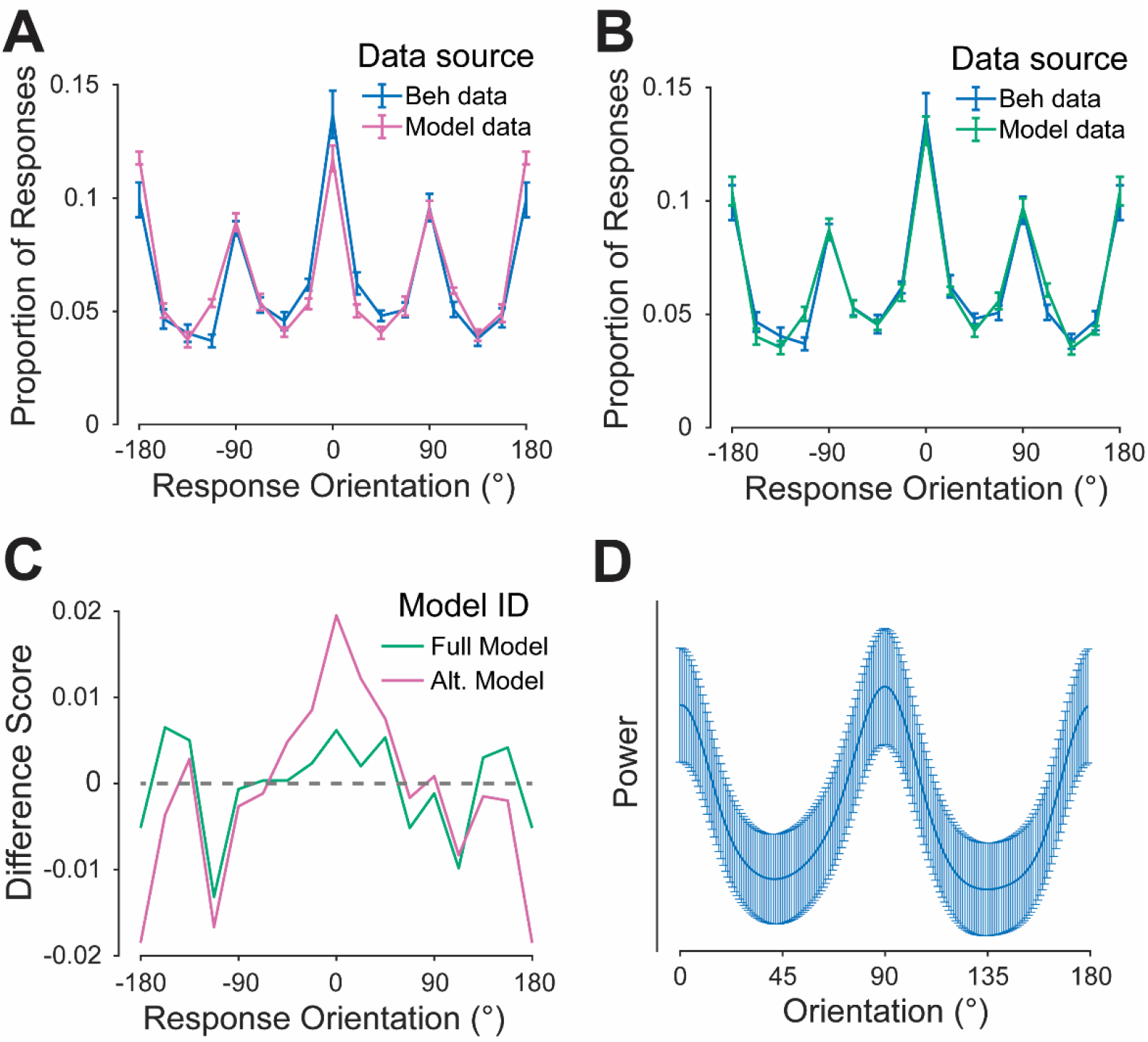
Perceptual inferences are predicted by natural image statistics. **A**) Proportion of response orientations (bin size = 22.5°) as measured from model and human observers in Experiment 1. The model judged the same targets as participants using an orientation heuristic. The model judged the same targets as participants using an orientation heuristic. **B**) The same behavioural data from Panel A, however model data in Panel B represent proportions of reported orientations yielded from a model that judged the same targets as participants using an orientation *and* lighting heuristic. **C**) Difference scores between model predictions and participant data, comparing the full model incorporating lighting and orientation heuristics (green) and the alternative model using an orientation heuristic only (pink). A difference score of 0 indicates the model predicted the same frequency of responses that participants displayed, positive values indicate the model predicted less responses than participants displayed, and negative values indicate the model predicted more responses than participants displayed. **D**) The mean distribution of contrast energy for each target when rotated to the orientation reported by the observer. Note this includes only the trials where participants responded with a >=90° response orientation. N = 10; error bars = ±1 SEM.

To understand how observers made their judgments in the absence of meaningful contextual information, we built a model observer that rotates each target so that the distribution of low-level features approximates the average distribution across many thousands of natural images. For each target we computed contrast energy across orientations, and calculated the circular shift required to minimise the difference between this distribution and the distribution of oriented contrast typically found in natural images, as reported previously (e.g., Hansen et al., 2003; Hansen & Essock, 2004; Harrison, 2022; see Methods). Note that this model is equivalent to asking what pattern of data would result from solely matching low-level target features to a prior acquired from averaging image statistics across many different scenes.

The output of this model is shown as the pink function in Figure 2A. Importantly, we did not fit the model to the observers’ data. Nonetheless, the model provides a very good approximation of their responses, with clear cardinal biases. However, the model underrepresents the frequency of responses around 0°, and overrepresents the frequency of responses around ±180°. This model error arises because oriented contrast energy is phase invariant, and so the orientation distribution is very similar for an upright or inverted image. We therefore added a second stage of the model that, after rotating the image to align its oriented contrast distribution to the prior, determines whether the target needs to be rotated a further 180°. This stage involved using a broadscale filter to estimate the lighting direction, which we fit to observers’ data with a single free parameter. The output of this full model is shown in Figure 2B, and now correctly estimates the frequency of responses for all orientations. Indeed, the two-stage model minimises the error in the overall model fit (Fig. 2C).

The results for Experiment 1 suggest that observers judge the appropriate appearance of an image patch by matching a relatively simple set of low-level image statistics to an average of these statistics over many images. If this is the case, observers’ responses should reflect the statistics of natural images even when their responses are highly inaccurate. We tested this prediction using reverse correlation to analyse the distribution of oriented structure for trials in which observers’ absolute response orientation was at least 90° from the ground truth orientation. For such trials, we computed oriented contrast energy for each target after rotating the target to the orientation reported by the observer. The mean distribution of contrast energy is shown in Figure 2D. There are clear peaks and troughs at cardinal and oblique orientations, respectively, aligning closely with contrast energy distributions in nature. Taken together, our model and reverse correlation analyses suggest that observers’ perceptual inferences depend on their prior expectations for the statistics of natural images.

### Contextual information enhances perceptual inferences

In Experiment 2, we next investigated the extent to which participants integrate context-specific information when inferring the upright orientation of novel target patches. Participants completed two blocks of trials in which we manipulated contextual information. In the “no-cue” block, the trial design was identical to that used in Experiment 1 above. In the “mixed-cue” block, participants performed the same task, but the target was preceded by either a contextual cue or a pink noise cue (see Fig. 3A). Contextual cues were the surrounding image from which a given target was drawn, with the target cropped out. By providing target-relevant contextual information, we directly investigated how observers weigh immediate contextual information relative to their longer-term priors. Furthermore, the use of a blocking design (Fig. 3B) allowed investigation into the temporal nature of contextual impacts on perceptual decisions. By interleaving contextual cues with noise cues in the mixed-cue block, we could further assess the degree to which contextual information carries- over from one trial to the next.

**Figure 3.**
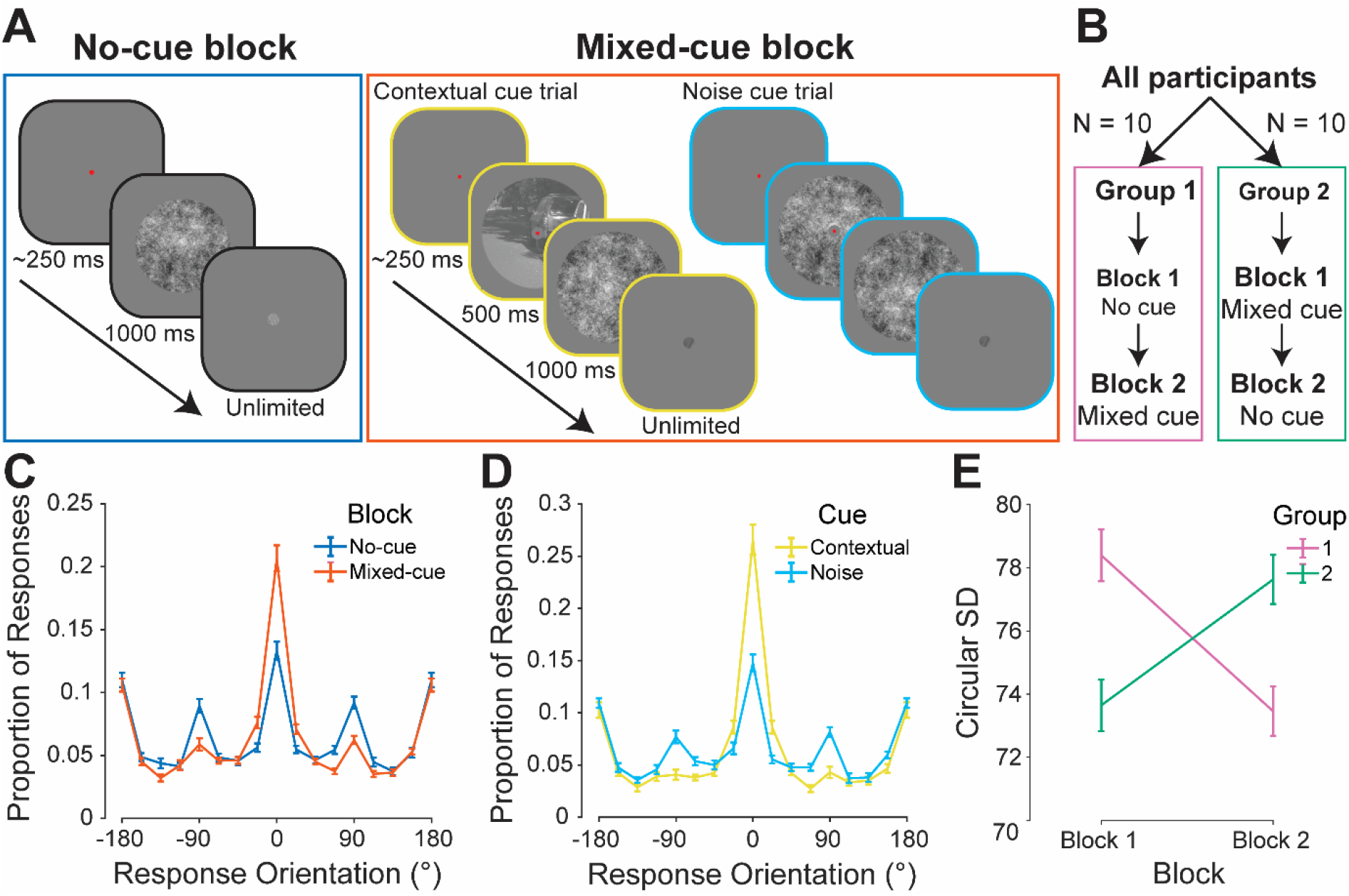
Contextual information enhances perceptual inferences above long-term priors. **A**) Example trial schematics for both tasks (no-cue and mixed-cue) in Experiment 2. **B**) Depiction of the mixed design of Experiment 2. Participants were split into two groups of 10. Each group did both tasks, but in different orders to eliminate the impact of potential practice effects on result interpretations. **C**) Comparison of mean responses for the no-cue (blue) and mixed-cue (orange) conditions. **D**) Comparison of performance for the two cue types *within* the mixed-cue condition: contextual cues (yellow) and pink noise cues (blue). **E**) Comparison of response variability for the two groups depicted in panel B: Group 1 (no-cue condition first; pink) and Group 2 (mixed-cue condition first; green). The x- axis indicates the block (a temporal category, indicating the order in which the two conditions were undertaken). The y-axis indicates the circular standard deviation of response orientations. N = 20; error bars: ±1 SEM.

The distributions of response orientations for the no-cue and mixed-cue conditions are shown in Figure 3C. The proportion of responses close to the objective upright (i.e., 0°) is almost two-fold greater in the mixed-cue condition than the no-cue condition, showing a clear facilitation of judgments when targets were preceded by a cue. Indeed, observers’ circular standard deviation was significantly lower in the mixed-cue condition (*M* = 73.55, *SD* = 3.52) than the no-cue condition (*M* = 78.01, *SD* = 1.58; BF_10_ = 42244.747; see Fig. 3C). By including contextual cues and noise cues in the mixed-cue condition, we further found that the changes in performance in the mixed-cue condition were driven by judgements only when contextual information was provided (Fig. 3D). These contextual effects cannot be accounted for by simple practice effects because there was no effect of block order on task performance (BF_10_ = 0.366; see Fig. 3B/E). Instead, the findings suggest participants’ judgements are influenced by transient contextual information, enhancing such judgements above the use of long-term priors alone. However, when contextual information is removed, observers again rely on priors to make their judgments about novel stimuli.

### Contextual benefits arise rapidly, but are image-specific

The results of Experiment 2 revealed that observers’ perceptual inferences involve a trade-off between their prior expectations and transient contextual information. Recent evidence suggests that contextual information is incorporated rapidly in the initial feedforward processing of an image (Neri, 2017). To determine when contextual effects emerge in our paradigm, we conducted Experiment 3 in which the duration of the contextual cue was manipulated: cues were presented for either 0.125, 0.25, 0.5, 1.0, 2.0 seconds, or not at all. This allowed us to investigate the time course over which contextual information is integrated.

The distributions of response orientations as a function of cue duration are shown in Figure 4A. The proportion of inverted responses is similar across cue durations, most likely because some targets are approximately symmetric across the horizontal axis (e.g., the bricks in Figure 1A). Nonetheless, the positive relationship between cue duration and proportion of responses centred on 0° demonstrates the increasing influence context-specific evidence with exposure. We summarise the data from each condition using circular standard deviation in Figure 4B, which shows an inverse relationship between response variability and cue duration, and reveals that contextual influences emerge rapidly, consistent with previous findings (Neri, 2017). We fit a hinged line to these data, which further reveals that contextual benefits accrue with increasing presentation times but then plateau after approximately 0.5 seconds, suggesting a maximal benefit of contextual exposure (Fig. 4B, hinged line). Hence, while contextual benefits appear to arise rapidly, their influences strengthen only up to a particular exposure time, after which additional benefits become negligible.

**Figure 4.**
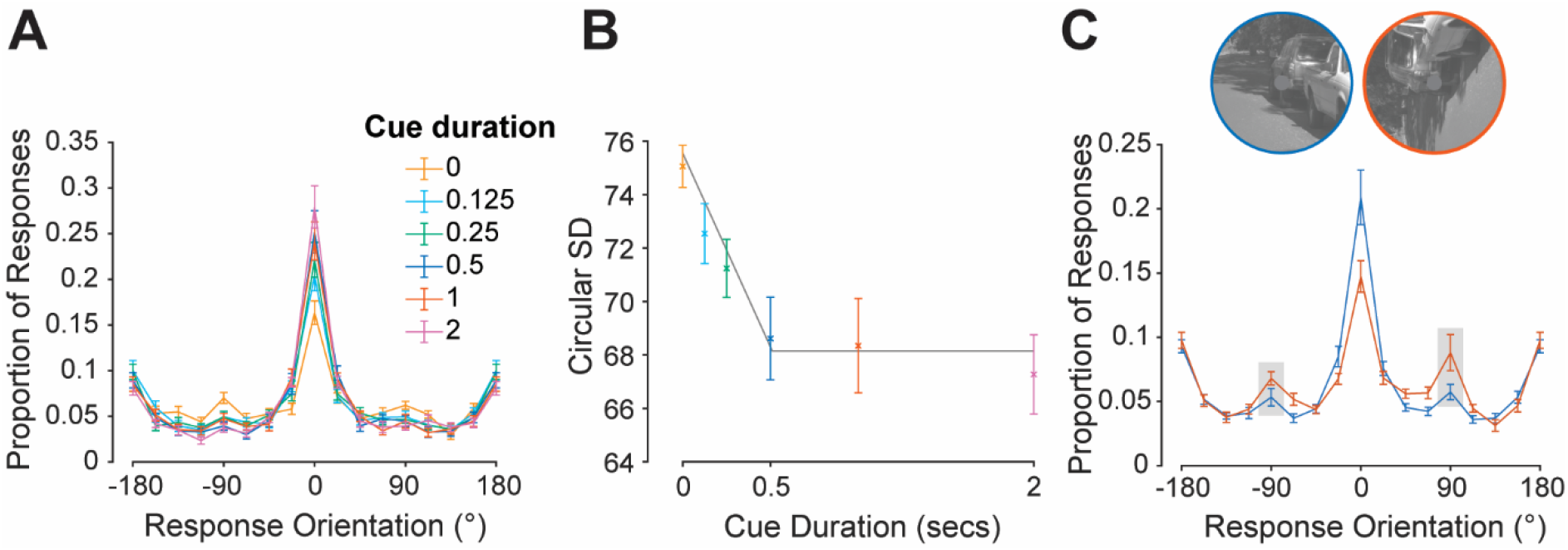
The timing and orientation of the cues influence contextual benefits. **A**) Results of Experiment 3, comparing mean response proportions for the six cue presentation time conditions (different conditions depicted by different colours). **B**) Comparison of response variability for the six cue durations (x-axis) as measured by circular standard deviation. A hinged line has been fit to the data to illustrate the point of maximal contextual benefit (grey). **C**) Results of Experiment 4, comparing performance in the upright-cue condition (blue) and rotated-cue (orange) condition, specifically for trials where contextual cues were given. Given that rotated cues were presented at +90° and -90°, responses for - 90° trials have been reverse-coded such that any expected biases away from a 0° response orientation due to rotated cues can be expected in the same direction (i.e., towards 90°). For ease of interpretation, data points of interest have been highlighted by grey boxes. We see a biasing of responses towards the presented cue orientation, with a greater proportion of 90° response orientations in the rotated cue condition compared to the upright cue condition, revealing a biasing effect of the cues’ low-level features. N = 20 per experiment; error bars: ±1 SEM.

### Contextual benefits arise regardless of relevance

Experiments 2 and 3 elucidated a strong influence of relevant contextual information on participants’ responses. However, it remains unclear whether participants utilise and become biased by contextual information because it is directly relevant to the task, or simply because observers automatically use any information regardless of its relevance. We investigated this in Experiment 4, presenting participants with upright contextual cues as well as cues rotated by ±90°. Importantly, participants were explicitly told whether cues were going to be upright or rotated in separate blocks and were instructed to rotate the target to be upright (i.e., not necessarily aligned with the contextual cue). In doing so, participants were actively discouraged from using rotated cues, allowing us to investigate whether participants still become biased by these less relevant cues.

We first standardised responses in the ±90° rotated-cue conditions by reverse-coding responses for trials where cues were rotated by -90°. As shown by the highlighted regions of the distributions in Figure 4C, responses were biased toward 90° in the rotated-cue condition compared to in the upright cue condition. We also observed an increase in responses in the opposite direction, i.e., toward -90°. This is consistent with participants being influenced by the rotated cue and inverting their response on a subset of trials. The presence of such biasing by unhelpful rotated cues suggests perceptual inferences cannot be made entirely independently of contextual information, even if that information is explicitly known to be irrelevant.

## Discussion

We investigated the contributions of long-term priors and immediate contextual information to perceptual inferences. Observers performed a task in which they rotated a target stimulus to its upright orientation, which required them to match the image features to some internal representation. Across four main experiments, we found that such perceptual inferences follow from priors for natural image statistics, and that the availability of transient contextual information enhances such inferences above the use of long-term priors alone.

### Perceptual inferences from natural image statistics

We found converging evidence that participants use priors for low-level image structure when interpreting isolated naturalistic image regions. In Experiment 1, participants’ inferences were explained by a model that matches targets to the distribution of orientation contrast seen in the natural world on average. Our reverse correlation analyses revealed that even large errors were aligned with the statistics of natural images on average (Fig. 2D). Therefore, our results demonstrate that judgments about the appearance of naturalistic images depend on priors accumulated over many scenes over a relatively long period of time.

While the tendency for observers’ responses to be biased toward cardinal orientations is similar to the oblique effect (Appelle, 1972; Berkley et al., 1975; Campbell et al., 1966; Dakin, 2001; Dakin et al., 2009; Dakin & Watt, 1997; de Gardelle et al., 2010; Emsley, 1925; Girshick et al., 2011; Pratte et al., 2016; Westheimer & Beard, 1998), our results cannot be explained by an oblique effect alone. Oriented contrast energy can only elucidate the horizontal and vertical axes of an image, not which half of the image should be at the top. Our full model therefore included a free parameter that estimated observers’ preferred lighting direction, analogous to the well-known light-from-above prior (Brewster, 1826; Metzger, 1936; Murray, 2013; Ramachandran, 1988), and aligned with behavioural responses. Hence, our results show that a relatively simple set of priors derived from natural image statistics is sufficient to drive heterogeneous patterns of perceptual inferences.

Despite the abstract appearance of many targets, observers’ modal response was accurate in all experiments. We designed targets so that there was no information that could unambiguously inform participants’ responses. We nonetheless ruled out any meaningful contribution of high-level structure in two control experiments: these experiments showed that the targets’ size and image content precluded observers’ use of high-level or semantic image content to inform their judgments (see Supplemental Materials). That observers’ modal response was accurate therefore reveals that low-level features provide sufficient information to interpret complex stimuli, such as natural image regions.

### Contextual information is assimilated rapidly

Our study demonstrates that perceptual inferences are guided by contextual information on the sorts of timescales that are behaviourally relevant. Such shifts in biases of perceptual judgements have been demonstrated for low-level features such as orientation (Lorenc et al., 2018; Rademaker et al., 2015; Taylor & Bays, 2018) and lighting (Adams et al., 2004; Morgenstern et al., 2011; Series & Seitz, 2013). Contextual influences have also been observed with high-level structures, with superior object detection and recognition when objects are embedded within contextually relevant scenes relative to contextually incongruent scenes or in isolation (Bar, 2004; Bar & Ullman, 1993; Biederman et al., 1982; Oliva & Torralba, 2007; Palmer, 1975). Importantly, we observe response bias shifts in response to contextual cues displayed for short durations. Such a time course is consistent with studies demonstrating our ability to rapidly process high-level information (e.g., scene categorisations and descriptions; Fei-Fei et al., 2007; VanRullen & Thorpe, 2001; Walther et al., 2009) and low-level information (e.g., feature averaging; Chong & Treisman, 2003; Parkes et al., 2001; Wolfe et al., 2011). Beyond supporting previous findings, our results suggest that the brain not only prioritises the rapid encoding and interpretation of complex stimuli, but that it is also able to utilise this information extremely effectively to inform judgements about subsequent stimuli.

### Contextual information is assimilated even when task-irrelevant

In Experiment 4, we found evidence suggesting that contextual information had an effect on perceptual inferences, even when participants were aware it was task-irrelevant. Previous literature has demonstrated detrimental impacts of incongruent context on visual detection/recognition tasks (Bar, 2004; Bar & Ullman, 1993; Biederman et al., 1982; Oliva & Torralba, 2007; Palmer, 1975). Such visual search tasks typically investigate the impact of irrelevant contextual information on an embedded target that cannot be separated from its contextual surroundings. Our study, however, allows us to investigate contextual influences under circumstances where participants are both aware of the unhelpfulness of the contextual information, and do not need to use it or explicitly engage with it to complete the task. The fact that we observe biasing towards irrelevant contextual information suggests that we interpret and incorporate contextual information regardless of relevance. Such a strategy should be beneficial under most circumstances, reflecting our experience in the real world where information is most commonly observed within its relevant context.

### Low-level image statistics and cognitive judgments

In the context of visual perception research, there has been substantial debate around the interactions between perceptual/bottom-up and cognitive/top-down processing (Firestone & Scholl, 2016; Pylyshyn, 1999). Relatively little is known about how such processes interact with one another in the context of interpreting natural scenes, with many investigations focusing on the contribution of high-level structures alone, and very few assessing the contributions of low-level perceptual processing. Such a focus on the contributions of high-level structure to cognitive tasks risks underestimating the potential contribution of low-level structure. Indeed, there is a precedent for such a notion, with evidence to suggest that systematic biases in visual working memory for simple stimuli can be accounted for by changes in basic image statistics (Taylor & Bays, 2018). Our results are in line with this finding: although our novel task involved what may be considered a relatively high-level task – to orient a random natural image patch to the subjective upright – observers’ performance can be captured by basic low-level image properties. Further, we have found clear contextual influences on such processes, with participants rapidly becoming biased by such information (whether relevant or irrelevant). Our results therefore provide insight into how interpretations of scene appearances follow from the most basic perceptual processes, which effectively assess a scene based on existing priors for such low-level information in tandem with available contextual information.

## Methods

### General task design

On each trial, participants were presented with one target patch, which was randomly oriented in the centre of the display (see Fig. 1A-B). Participants’ task was to infer the “upright” orientation of the target by rotating it using a mouse. Specifically, participants were instructed that they would see a series of “targets”, each cut out from a larger image, the “source image”, and presented at a random orientation. Participants were instructed to rotate these targets to their upright orientation. Performance was measured using the orientations reported by the participant for each trial relative to the objective upright orientation of the patch (see Fig. 1C).

### Participants

Numbers of participants differed across experiments, ranging from N=10-20. Participants had varying degrees of experience participating in psychophysical experiments, but all participants were naïve to the purpose of the experiments except for one participant in Experiment 1 (an author). Ethics approval was granted by the University of Queensland Medicine, Low & Negligible Risk Ethics Sub-Committee.

### Stimuli

Targets and contextual cues were generated in the same manner across all experiments unless otherwise specified. Digital natural images were taken from a database of high-resolution colour photos, cropped to 1080×1080 pixel regions (Burge & Geisler, 2011). For Experiments 1-4, target patches were circular patches cropped from the centre of the 1080×1080 images, subtending 2° of visual angle in diameter. In experiments where contextual cues were given, the 1080×1080 images that the target patches were cropped from were used. These were cropped to an annulus with a 27° outer diameter and a 2° inner diameter (i.e., cropping out the target patch). All stimuli were converted to greyscale using Matlab’s rgb2gray() function.

### Apparatus

Stimuli were displayed on a Dell Precision T1700 computer (running Windows 7 Enterprise) with the Psychophysics Toolbox (3.0.12; Brainard, 1997; Pelli, 1997) for MATLAB (R2015a). Stimuli were presented on a 24-inch Asus VG428QR 3D monitor with 1920 × 1080-pixel resolution and a refresh rate of 100 Hz. A gamma correction was applied to the display, assuming that gamma was 2.

### Experiment 1 Design

Participants (N=10; no exclusions) completed 600 trials. For each participant, 150 unique target patches were pseudorandomly selected from a bank of 9,361 potential targets. Participants performed the task of re-orienting a randomly oriented target patch to be “upright” on each trial with no additional contextual information given about the patches (see Fig. 1A-B). Prior to completing the experiment, participants did 20 practice trials. Trial order was randomised and sessions were split into six blocks of 100 trials with self-timed breaks in between.

In Experiment 1, four copies of each target patch were made, with each copy having a different level of white noise (0%, 2.5%, 5%, and 10%) applied through manipulating the RMS contrast of the image. This was an initial line of interest in the current study, however there was no significant effect of target noise on participants’ performance as measured by circular standard deviation (BF_10_ = 0.933). We therefore combined these conditions in the data shown in Figure 2 and the manipulation was not explored in further experiments.

### Experiment 2 design

Twenty-one participants completed 600 trials split into two blocks (see Fig. 3A). One dataset was not analysed because the participant did not finish the experiment. In one block, each participant performed the basic task of re-orienting 300 target patches to be “upright” (the “no-cue” block). In the other block, participants completed the same task, but a contextual cue (i.e., the surrounding source image the target was drawn from with the target cropped out) or a random pink noise patch was displayed for a duration of 500 ms prior to the target (the “mixed-cue” block). For the mixed-cue block, 150 target patches were pseudorandomly selected for each participant: each unique target patch was presented twice – once preceded by a pink noise patch (noise cue) and once preceded by the surrounding image from which the target was drawn (contextual cue). Participants completed 20 practice trials prior to completing each block, with practice trials implementing the same cue types as the blocks. Block order was counterbalanced across participants (see Fig. 3B). Trial order for each block was randomised and each block was split into three sets of 100 trials with self-timed breaks in between. On average, participants took 29 minutes to complete each block (excluding three participants, due to a duration recording error).

### Experiment 3 design

Participants (N=21; one excluded due to not finishing the experiment) completed 450 trials. For each participant, 75 unique target patches were pseudorandomly selected. Unique target patches were presented six times, preceded by the surrounding image from which the target was drawn – once for each presentation length: 0, 0.125, 0.25, 0.5, 1.0, and 2.0 seconds. Prior to completing the experiment, participants did 20 practice trials. Trial order was randomised and sessions were split into six blocks of 75 trials with self-timed breaks in between. On average, participants took 48 minutes to complete the task.

### Experiment 4 design

Participants (N=20; no exclusions) completed 600 trials split into two blocks. In one block, participants completed the base task, however before seeing the target patch participants were either shown a typical contextual cue or a random pink noise patch, with 150 pseudorandomly selected target patches for each participant (the “upright-cue” block; see Fig. 4C). The other block had an identical design, however contextual cues (when presented) were rotated by ±90° (the “rotated-cue” block; see Fig. 4C). Block order was counterbalanced across participants. Participants completed 20 practice trials prior to completing each block, with practice trials implementing the same cue types as the blocks. Each block was split into three sets of 100 trials with self-timed breaks in between. On average, participants took 31 minutes to complete each block.

### Control experiments

Targets were generated in a uniform manner across images in our database and were not screened for their content before being shown to participants. Therefore, in cases where participants were not given contextual information, the possibility remained that the targets themselves included sufficient high-level structure to unambiguously cue their objective upright orientation. The presence of such high-level information could potentially explain participants’ performance. For example, if participants happened to be shown a picture of a car, they would be expected to know from their semantic knowledge which way is upright. As such, two control experiments were conducted to investigate the potential contribution of high-level structure in the isolated target patches to participants’ performance. These experiments are summarised below and described in detail in the Supplemental Materials.

The first control experiment had participants categorise individual patches according to whether there was sufficient high-level information present to unambiguously indicate the “correct” upright orientation of the image (informative) or not (uninformative). This experiment found there was neither sufficient numbers of informative images shown to participants, nor consistently accurate responses made in response to informative images to account for levels of performance observed. Thus, informative image regions are unable to account for participants’ inference judgements of the upright appearance of naturalistic images.

The second control experiment involved the same trial structure and task as Experiment 1, but the size of the target patch was manipulated to include varying amounts of the source image. We found that as target size increased (and therefore the amount of high-level structure), so did performance. This pattern suggests that the decision to limit the target size was effective in eliciting task difficulty by limiting the amount of high-level information present. Together, our two control experiments revealed relevant high-level information content that disambiguated the perceptual task was almost entirely absent for our target stimuli and therefore cannot account for participants’ performance.

### Analyses

Where inferential statistics were performed, Bayesian analyses were implemented in JASP, using circular standard deviation (i.e., response orientation variance, where greater variance indicates poorer performance) as the dependent variable. For Experiment 3 response variability data (Fig. 4B), we fit a hinged line by finding the parameters that minimised the square error between each participant’s data and the model using MATLAB’s fminsearch() function.

### A “pretty good observer” model of upright inferences

We developed an observer model that estimates the upright orientation of a target patch by matching the target’s statistics with the anisotropic distribution of orientation energy found in nature (e.g., Hansen et al., 2003; Hansen & Essock, 2004; Harrison, 2022). We refer to this model here as a *pretty good* observer model, rather than an *ideal* observer model, because we only exploit orientation energy and ignore other statistical features that could further improve performance (e.g., conditional orientation statistics; Geisler et al., 2001). For a given target patch, we computed orientation energy in 180 equally spaced orientation bands, each of which covered all spatial frequencies. These operations were performed in the frequency domain; energy was the absolute of the Fourier-transformed target values. Oriented filters were also constructed in the frequency domain: filters were raised cosines with a bandwidth of 45°. Energy was summed within each orientation band, giving a distribution of energy across orientations. We then compared the target’s energy distribution to a prior derived from studies of natural images (e.g., Wei & Stocker, 2015):

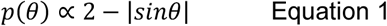

Where *p*(*θ*)is the probability of observing contrast energy with an orientation of *θ*, in radians. Whereas Equation 1 assumes equal prevalence of horizontal and vertical orientations, Hansen and colleagues noted there tends to be a horizontal bias (see also Harrison, 2022). We therefore modified Equation 1 by increasing the proportion of horizontal energy according to a von Mises function:

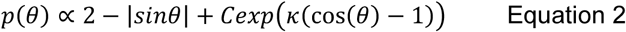

Where C is the strength of the horizontal bias, and *k* is the width of the von Mises function, which we set to 2.5. Small changes in *k* did not change the results. Before summing the distributions, we first normalised the von Mises function to have a peak of one.

The model selects the upright orientation of a target by computing the rotational offset that minimises the sum of the squared difference between the target patch’s energy and *p*(*θ*), performed using MATLAB’s fminsearch() function. Prior to this step, we further normalised the target’s energy distribution and *p*(*θ*)to both fall within the range 0 – 1. To avoid local minima, we fit the model with varying starting parameters and took the rotational offset at the global minimum from all fits. Note that this fitting is entirely independent of an observer’s response – we fit the target’s image statistics to the prior, but nonetheless approximate observers’ responses very closely (Figure 1A). However, as described in the Results, this model produces an equal proportion of 0° and ±180° responses, which is different than observers’ reports. We therefore added a second stage of the model that, after finding the best rotational offset, estimates lighting direction from a broadscale filter positioned at the centre of the target. The orientation of this filter matched the fitted offset. Crudely, this filter can be considered an estimate of the relative phase of a horizon running through the middle of the target. Depending on the polarity of the filter’s response, the image was either rotated a further 180°, or left as is. This step considerably improved the match between the model responses and observers’ data, as shown in Figures 1A-1C.

## Acknowledgements

We would like to thank Micaela Dear for her assistance with data collection.

The current study was funded by an ARC DECRA to WJH (DE190100136).

## Supplemental Materials

## Control Experiments

Targets were generated in a uniform manner across images in our database and were not screened for their content before being shown to participants. Therefore, in cases where participants were not given contextual information, the possibility remained that the targets themselves included sufficient high-level structure to unambiguously cue their objective upright orientation. The presence of such high-level information could potentially explain participants’ performance. For example, if participants happened to be shown a picture of a car, they would be expected to know from their semantic knowledge which way is upright. As such, two control experiments were conducted to investigate the potential contribution of high-level structure in the isolated target patches to participants’ performance.

### Participants

Numbers of participants differed across experiments, ranging from N=2-3. Participants had varying degrees of experience participating in psychophysical experiments, and all participants were naïve to the purpose of the experiments except for one participant in both experiments (an author). Ethics approval was granted by the University of Queensland Medicine, Low & Negligible Risk Ethics Sub-Committee. **Apparatus** Stimuli were generated on a Dell Precision T1700 computer (running Windows 7 Enterprise) with the Psychophysics Toolbox (3.0.12; Brainard, 1997; Pelli, 1997) for MATLAB (R2015a). Stimuli were presented on a 24-inch Asus VG428QR 3D monitor with 1920 × 1080-pixel resolution and a refresh rate of 100 Hz. A gamma correction was applied to the display, assuming that gamma was 2.

### Control Experiment 1

#### Stimuli

Stimuli were made up of targets used in Experiment 1 and 2, as well as an additional unreported experiment that implemented the same design as Experiment 2 but did not involve a no-cue block. In total, there were 7176 unique targets shown to participants across these three experiments. Briefly, these targets were digital natural images taken from a database of high-resolution photos (Burge & Geisler, 2011) cropped to subtend 2° of visual angle in diameter and converted to greyscale (see Methods for more detail).

#### Design

Participants (*N* = 2; one author) completed 7176 trials. On each trial, participants were shown a target in its upright position. For each target, participants judged whether there was sufficient high-level information present to unambiguously indicate the “correct” upright orientation of the image (“informative”) or not (“uninformative). For example, patches that contained identifiable objects such as cars or signs were classified as informative. Categorisations were made using the 0 (uninformative) and 1 (informative) keys on a keyboard. Targets were presented until a response was made, and participants had the ability to backtrack using the ‘Backspace’ key. Participants completed the experiment across self-selected block lengths.

#### Analyses

Results presented here are based on Rater 2’s (non-author) data, who categorised more images as “informative” (295 of 7176) than Rater 1 (66 of 7176; an author). Results are based on Rater 2 as we want to be conservative in attributing effects to low-level features relative to high-level features in our targets. Hence, by basing analyses on Rater 2’s more liberal informative categorisations, we give informative images the greatest chance of explaining the observed patterns of data across experiments.

#### Results

In Experiment 1, there was a sub-sample of image patches categorised as informative (58 of 1400 unique targets across 10 participants; 4%). On average, 6.20 (*SEM* = 0.63) unique informative images were shown to each participant, accounting to 4% of the total number of unique targets seen. Across participants, we see a range of response orientations when informative images are shown (Fig. S1A), with informative images responded to accurately on 29% of trials when they are shown. Hence, even when presented with informative image regions, participants do not necessarily perform optimally, diminishing the explanatory power of informative images accounting for accurate responses. Indeed, on average, accurate responses attributed to informative images account for just 1% of participants’ responses (Fig. S1A).

**Figure S1.**
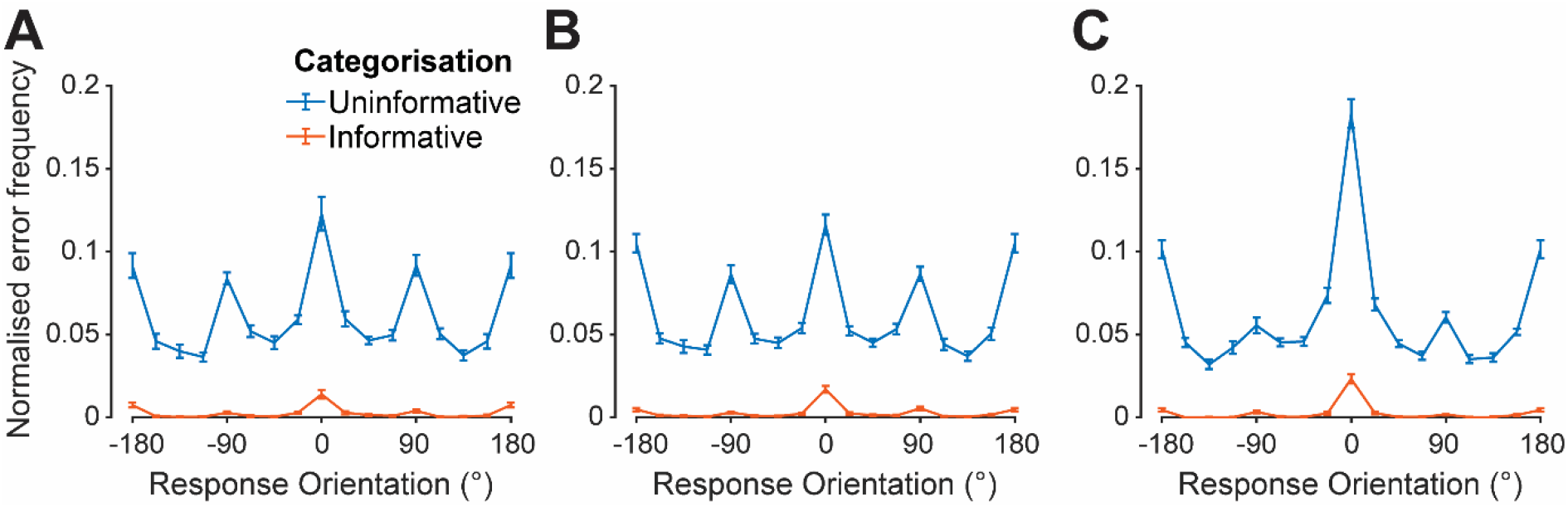
The contribution of informative images to participants’ responses. **A**) Proportion of responses made in Experiment 1, split by responses made to targets categorised as informative (orange) and uninformative (blue). **B**) Proportion of responses made in the no-cue block of Experiment 2, split by responses made to targets categorised as informative (orange) and uninformative (blue). **C**) Proportion of responses made in the mixed-cue block of Experiment 2, split by responses made to targets categorised as informative (orange) and uninformative (blue). N = 10 (A) & 20 (B/C); error bars: ±1 SEM.

This was reflected in Experiment 2 stimuli. In the no-cue block, 190 (4%) of the 4510 unique targets used were rated as informative. On average, 13.5 (*SEM* = 0.92) unique informative images were shown to each participant, accounting for 5% of the total number of unique targets seen. Again, a range of response orientations for informative images was observed (Fig. S1B), with informative images responded to accurately on 34% of trials when they are shown. Similarly, on average, accurate responses attributed to informative images account for just 1% of participants’ responses (Fig. S1B).

Similarly, in the mixed-cue block,107 (4%) of the 2609 unique targets were rated as informative. On average, 6.3 (*SEM* = 0.54) unique informative images were shown to each participant, accounting for 4% of the total number of unique targets seen. Again, a range of response orientations for informative images was observed (Fig. S1C), with informative images responded to accurately on 50% of trials when they are shown. On average, accurate responses attributed to informative images account for just 2% of participants’ responses (Fig. S1C).

Taken together, these results demonstrate there was neither sufficient numbers of informative images shown to participants, nor consistently accurate responses made in response to informative images to account for levels of performance observed. Thus, informative image regions are unable to account for participants’ inference judgements of the upright appearance of naturalistic images.

### Control Experiment 2

#### Stimuli

Targets were digital natural images were taken from a database of high-resolution photos (Burge & Geisler, 2011). For each participant, 120 unique images were selected from the database. Each unique image had five copies generated, each cropped to circular patches of different sizes (1, 2, 4, 8, and 16°), such that different target sizes included varying amounts of the source image (Fig. S2A). All stimuli were converted to greyscale.

#### Design

Participants (*N* = 3; one author) completed five blocks of 120 trials, where each block corresponded to one target size (i.e., 1, 2, 4, 8, or 16°). Block order was randomised for each participant. The task was identical to Experiment 1, requiring participants to rotate a randomly oriented target patch to be perceptually upright.

#### Results

As anticipated, when the aera of the original image presented increased beyond that used in Experiments 1-4 (2° of visual angle), performance steadily increased, supported by large increases in the number of responses centred on a response orientation of 0° (Fig. S2B) and large decreases in response variability as measured by circular standard deviation (Fig. S2C). Improved performance with larger patch sizes suggests that providing larger patches, and therefore more high-level structure, makes the patches more informative and decreases the difficulty of the task.

**Figure S2.**
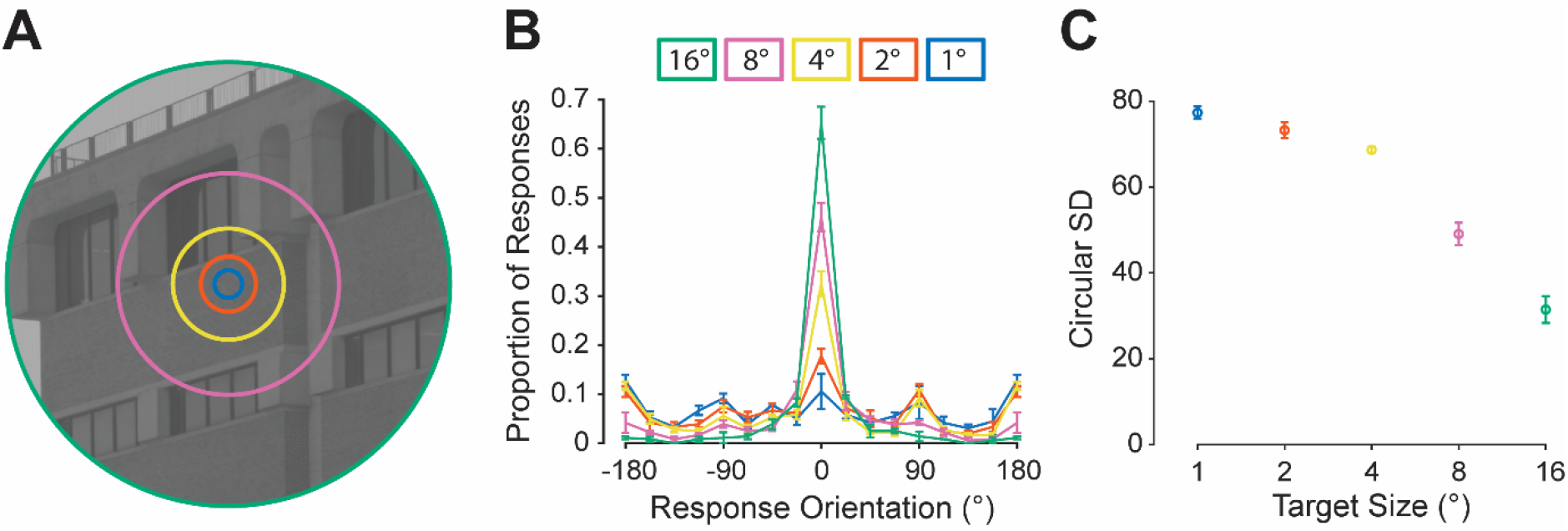
The impact of image size on perceptual inferences for naturalistic images. **A**) Depiction of target sizes used (1, 2, 4, 8, and 16°; smaller than actual size, but to scale relative to one another). **B**) Comparison of mean response proportions for the five target size conditions (different conditions depicted by different colours). **C**) Comparison of response variability for the five target sizes (x-axis) as measured by circular standard deviation. N = 3; error bars: ±1 SEM.

#### Overview

Overall, results from our informativeness and patch size control experiments suggest that limiting the targets to a windowed patch, particularly at our chosen patch size, was effective at removing much of the high-level structure present. Taken together with the four main experiments, the results presented suggest that priors for statistical regularities of low-level features in nature are sufficient to make informed interpretations of isolated naturalistic stimuli.

